# Bacterial derived vitamin B12 enhances predatory behaviors in nematodes

**DOI:** 10.1101/797803

**Authors:** Nermin Akduman, James W. Lightfoot, Waltraud Röseler, Hanh Witte, Wen-Sui Lo, Christian Rödelsperger, Ralf J. Sommer

**Affiliations:** Department for Evolutionary Biology, Max Planck Institute for Developmental Biology, Max Planck Ring 9, 72076 Tübingen Germany

**Keywords:** Microbiome, microbiota, diet, surplus killing, metabolism, development, *Pristionchus pacificus*, *Caenorhabditis elegans*

## Abstract

The microbiome is known to affect host development, metabolism and immunity, however, its impact on behaviors is only beginning to be understood. Here, we investigate how bacteria modulate complex behaviors in the nematode model organism *Pristionchus pacificus. P. pacificus* is a predator feeding on the larvae of other nematodes including *Caenorhabditis elegans*. Growing *P. pacificus* on different bacteria and testing their ability to kill *C. elegans* reveals drastic differences in killing efficiencies with a *Novosphingobium* species showing the strongest enhancement. Strikingly, increased killing was not accompanied by an increase in feeding, a phenomenon known as surplus-killing whereby predators kill more prey than necessary for sustenance. RNA-seq revealed widespread metabolic rewiring upon exposure to *Novosphingobium*, which facilitated the screening for bacterial mutants leading to an altered transcriptional response. This identified bacterial derived vitamin B12 as a major micronutrient enhancing predatory behaviors. Vitamin B12 is an essential cofactor for detoxification and metabolite biosynthesis and has previously been shown to accelerate development in *C. elegans*. In *P. pacificus* vitamin B12 supplementation amplified, whereas mutants in vitamin B12-dependent pathways reduced surplus-killing. This demonstrates that bacterial vitamin B12 affects complex behaviors and thus establishes a connection between microbial diet and the nervous system.

---

The microbiome is considered a fundamental aspect of a host’s biology and is known to provide developmental cues, influence metabolism and alter immunity^1-3^. However, the microbiome constitutes a complex network of microorganisms and disentangling specific interactions and effects at a mechanistic level is challenging. Bacterial-feeding nematodes constitute a highly attractive system to study the influence of the microbiome because specific interactions can be investigated in monoxenic cultures where the microbiome and diet are indistinguishable from one another and easily controlled. To study the effect of bacteria on behavior we investigate the nematode model organism *Pristionchus pacificus* that exhibits a particular complex behavior unknown from *C. elegans*. In general, *P. pacificus* is an omnivorous nematode that can grow on bacteria, fungi and it can predate on other nematodes^4-6^. Predation is dependent on morphological and behavioral novelties, involving the formation of teeth-like denticles and a self-recognition mechanism^7-10^. The ability to form teeth-like denticles is an example of developmental plasticity with two discrete mouth-forms^11^. The stenostomatous morph has a single blunt tooth, whereas the eurystomatous morph has two large teeth with only the latter capable of predation (Fig. 1A and B)^7^. Predation may confer a selective advantage in certain environmental settings with previous studies indicating that different culture conditions, including microbial diet, are able to modulate the ratio of the two mouth forms^12,13^. Furthermore, *P. pacificus* predation under laboratory conditions is also an example of a phenomenon known as surplus-killing behavior^6^. Surplus-killing is a well-documented complex behavior observed in many predators across the animal kingdom, in which more prey are killed than nutritionally required^14-22^. Theoretical and experimental studies considered surplus-killing a potentially context-dependent, adaptive foraging strategy or alternatively, a context-general syndrome of high aggression^15,17,20-23^. However, the full impact of diet on killing and predation is currently poorly understood.

**Figure 1.**
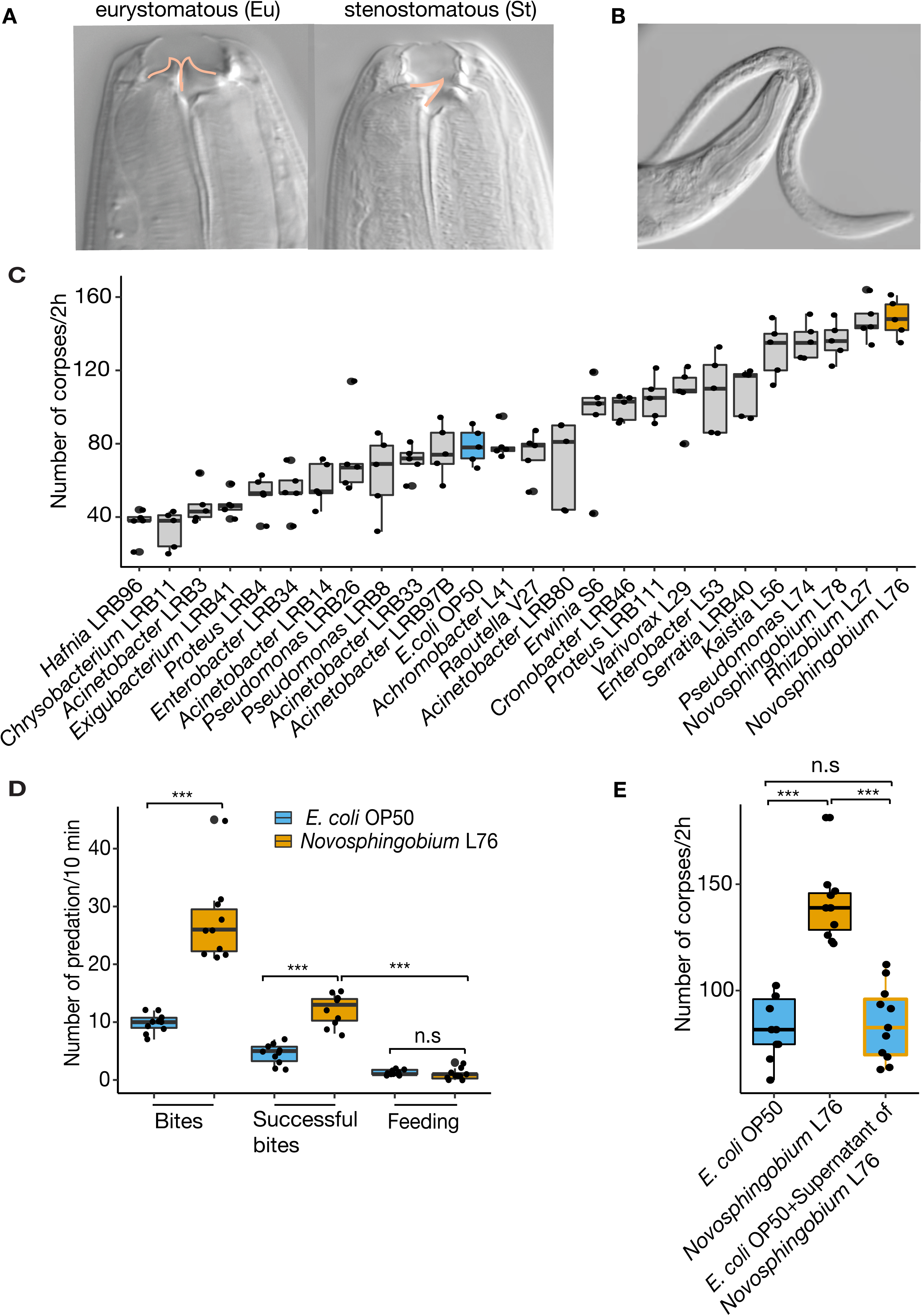
Bacterial diet modulates killing behavior in *P. pacificus*. (**A**) Eurystomatous (Eu) and stenostomatous (St) mouth forms. Eu worms are capable of predation and have a wide mouth with two teeth, while St worms feed on bacteria and have a narrower mouth with one tooth. (**B**) A predatory *P. pacificus* adult biting a *C. elegans* larvae. (**C**) Corpse assay of *P. pacificus* predators fed upon *C. elegans* larvae following growth on a variety of ecologically associated bacteria; five predators are fed prey for two hours for each assay. *N* = 5 replicates for each assay. (**D**) Bite assay after growth on either an *E. coli* OP50 or *Novosphingobium* L76 diet to assess the effect on *P. pacificus* surplus-killing behavior. Numbers of bites, successful bites and feeding was quantified during a 10 min interval while fed upon *C. elegans* larvae. (**E**) A corpse assay of *P. pacificus* fed with *E. coli* OP50, *Novosphingobium* L76 or of *E. coli* OP50 with *Novosphingobium* L76 supernatant. N=10 replicates for each assay for (D) and (E).

Therefore, we tested the effect of 25 different bacteria recently isolated from *Pristionchus*-associated environments^24^ on various predation associated traits. Specifically, we grew *P. pacificus* for several generations on monoxenic cultures and investigated the effect on mouth form ratio, pharyngeal pumping, and killing behavior by comparing them to standard laboratory cultures grown on *Escherichia coli* OP50. While diet had a limited effect on mouth form ratios and pharyngeal pumping, we found up to a four-fold difference in killing efficiency depending on microbial diet (Fig. 1C, fig. S1A and B). The strongest effect on killing efficiency was observed when *P. pacificus* was fed upon three alpha-proteobacteria of the genera *Novosphingobium* and *Rhizobium*, resulting in up to 160 corpses of dead prey in standardized corpse assays (Fig. 1C). We therefore focused on one bacterium of this group, *Novosphingobium* L76.

Stronger killing efficiency translated into higher rates of surplus-killing. Specifically, we performed bite assays to observe individual predators for 10 minutes to distinguish specific predatory events including biting, successful biting that results in penetration of the cuticle, and feeding on prey larvae (see Method section for exact description of terms). When grown on *E. coli* OP50, *P. pacificus* only kills 50% of its prey after biting, and subsequent feeding was only observed in roughly 10% of all cases (Fig. 1D, Movie S1). Using *Novosphingobium* L76, we found that the number of *P. pacificus* bites and successful biting events indeed doubled relative to *E. coli* OP50 grown predators (Fig. 1D). However, we found no increase in feeding on the dead prey (Fig. 1D). Instead, predators rapidly moved over agar plates searching for new prey items. Thus, a *Novosphingobium* diet enhances predation and surplus-killing.

Next, we established the necessary bacterial exposure time required to influence predatory behavior and additionally, wanted to know whether the increase in killing was mediated by factors secreted by the bacteria or solely by their ingestion. Only a limited exposure to a diet of *Novoshingobium* L76 during development was sufficient for *P. pacificus* nematodes to exhibit increased predatory behavior, however, *Novosphingobium* L76 culture supernatants alone were unable to recapitulate this effect (Fig. 1E, fig. S1C). In contrast, when *Novosphingobium* was diluted with *E. coli* OP50, the effect still persisted suggesting that the response to *Novosphingobium* L76 is unlikely due to differences in caloric intake (fig. S1D). Instead, the behavioral change is likely a result of physiological alterations caused by the different nutritional composition of *Novosphingobium* L76. Therefore, we analyzed the transcriptomic response of young *P. pacificus* adults grown on *Novosphingobium* in comparison with *E. coli*. We identified a total of 2,677 (9%) genes with significant differential expression (FDR corrected P-value < 0.05) between the two bacterial diets (Table S1). Most strikingly, more than half of all genes that are predicted to be involved in fatty acid metabolism are significantly differentially expressed between the two diets (Fig. 2A and B).

**Figure 2.**
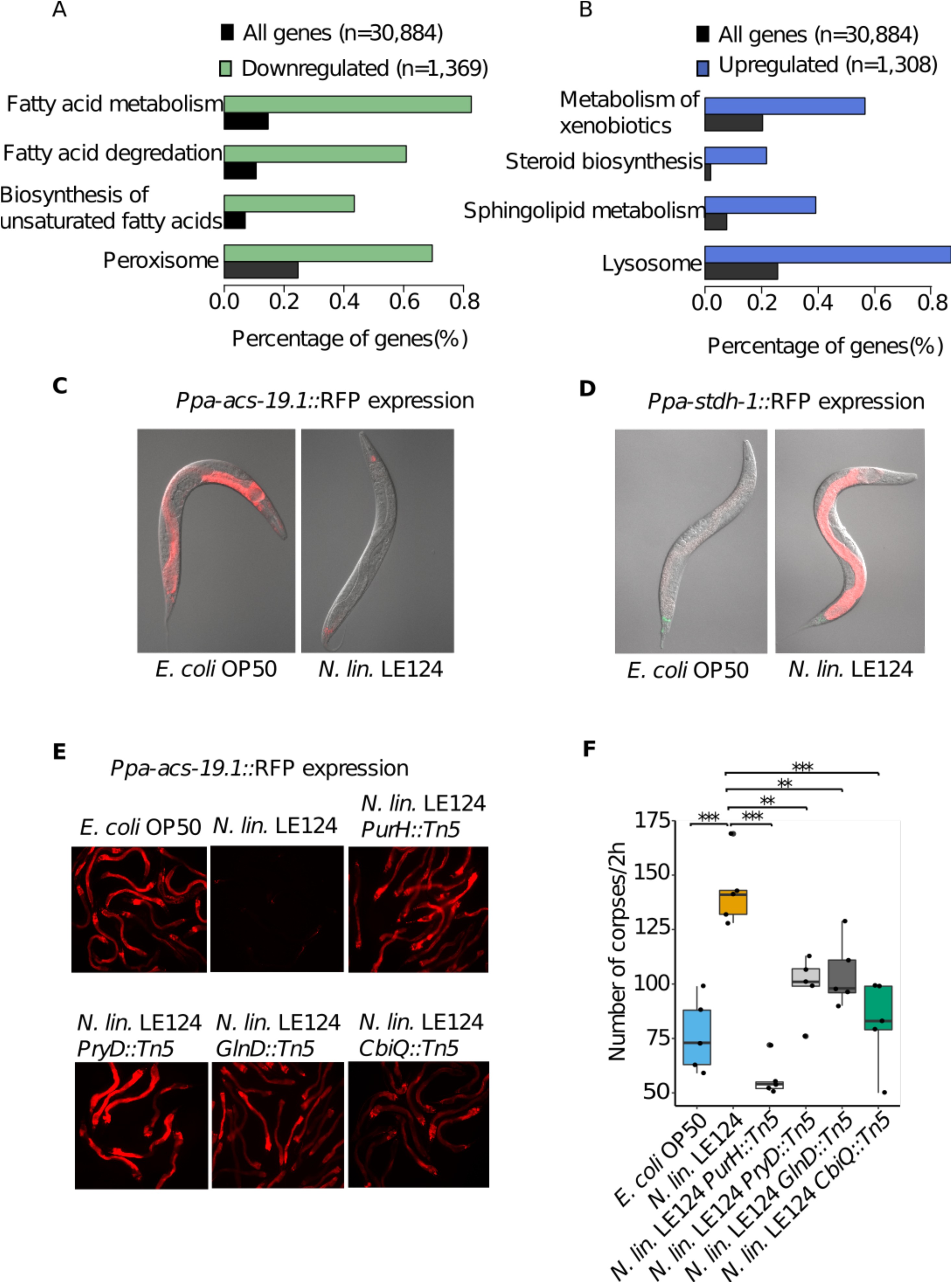
Bacterial diet influences gene expression in *P. pacificus*. (**A)** RNA-seq analysis of *P. pacificus* in response to a diet of *Novosphingobium* L76 compared to *E. coli* OP50. The pathways with most significant enrichment (FDR-corrected *P<10*^*-5*^) in downregulated and (**B**) upregulated genes are shown. (**C**) The dietary sensor *Ppa-acs-19.1*::RFP is highly expressed in ventral gland, hypodermal and intestinal cells following an *E. coli* OP50 diet, while a *Novosphingobium* L76 diet induces expression only in ventral gland cells. The co-injection marker *Ppa-egl- 20*::RFP is expressed in the tail. (**D**) *Ppa-stdh-1:*:RFP is expressed in the intestinal and hypodermal cells with expression strongly upregulated on *Novosphingobium* L76 diet compared to an *E. coli* OP50 diet. (**E**) Expression of *Ppa-acs-19.1*::RFP dietary sensor after feeding on *N. lin*. LE124 transposon mutants with mutations in vitamin B12 (*N. lin*. LE124 *CbiQ::Tn5)*, purine (*N. lin*. LE124 *PurH::Tn5*), pyrimidine biosynthesis (*N. lin*. LE124 *PryD::Tn5*) and nitrogen metabolism (*N. lin*. LE124 *GlnD::Tn5*). Mutants increase the expression of the dietary sensor in comparison to a *N. lin*. LE124 wild-type diet. (**F**) Corpse assay of *P. pacificus* after feeding on various *N. lin*. LE124 mutants. There is decreased killing efficiency compared to a *N. lin*. LE124 wild type diet. N=10 replicates for each assay.

To study the mechanisms by which *Novosphingobium* alters fatty acid metabolism and induces behavioral changes in the nematode, we used an unbiased bacterial mutagenesis approach. We replaced *Novosphingobium* L76 with *Novosphingobium lindaniclasticum* LE124 (*N. lin*. LE124 thereafter), as the latter can easily be manipulated by transposon mutagenesis, has an available genome^25^, and induces similar behavioral effects in *P. pacificus* (fig. S1E). Additionally, to detect any physiological changes in *P. pacificus* caused by mutations in the bacteria, two dietary sensors were generated using *P. pacificus* fatty acid metabolism genes that showed differential expression on different bacteria (Fig 2A and B). Specifically, we used homologs of the acyl-CoA synthetase enzyme *Ppa-acs-19.1*, which was upregulated on *E. coli* OP50 and downregulated on *Novosphingobium*, as well as the short-chain dehydrogenase reductase enzyme *Ppa-stdh-1*, which has the opposite expression profile (Fig. 2C and D, fig. S2). Both reporter lines confirmed the differential expression that was detected by RNA-seq with *Ppa-acs-19.1* being expressed nearly exclusively on *E. coli*, whereas *Ppa-stdh-1* is expressed highly on *Novosphingobium* but only minimally on *E. coli* OP50 (Fig. 2C and D). Subsequently, we used these dietary sensors to screen for bacterial mutants that fail to differentially regulate these genes. From a library of 4,320 *N. lin*. LE124 mutants, three affected the expression of *Ppa-stdh-1* and 21 altered the expression of *Ppa-acs-19.1*. Whole genome sequencing of these bacterial mutants identified transposon insertions in genes corresponding to four biological pathways: purine and pyrimidine metabolism, nitrogen metabolism, and vitamin B12 (Fig. 2E; fig. S2C, key resources table). Importantly, in mutants of all four pathways, the change of transcriptomic response coincided with a reduction in predatory behavior including surplus-killing relative to wild-type *N. lin*. LE124 (Fig. 2F, fig. S2D and E). Thus, the dietary sensor allows the identification of factors regulating complex behavioral traits.

Vitamin B12 has been shown to be a crucial co-factor involved in growth, development and behavior in several animals, including mice and human^26^. Therefore, we focus on vitamin B12, which was recently also found to affect growth and development of *C. elegans*^27^, whereas nothing is known about vitamin B12 affecting *C. elegans* behavior. We first analyzed if vitamin B12 supplementation was sufficient to affect the expression of the *Ppa-acs-19.1* sensor and determined the required concentration for this. Supplementation of an *E. coli* diet with 500nM vitamin B12 resulted in the absence of *Ppa-acs-19.1* expression with no adverse effects to the health of wild-type animals (fig. S3A). Additionally, this vitamin B12 concentration abolished *Ppa-acs-19.1* expression on *N.lin*.LE124 *CbiQ::Tn5* mutants (fig. S3B). Subsequently, we analyzed if this supplementation was also sufficient to enhance the predatory behaviors. Indeed, supplementation with 500nM vitamin B12 rescued the vitamin B12-deficient *N. lin*. LE124 *CbiQ* mutant and similarly, increased surplus-killing behavior on an *E. coli* diet (Fig. 3A and B). These results demonstrate that vitamin B12 is an important micronutrient involved in complex behaviors in nematodes.

**Figure 3.**
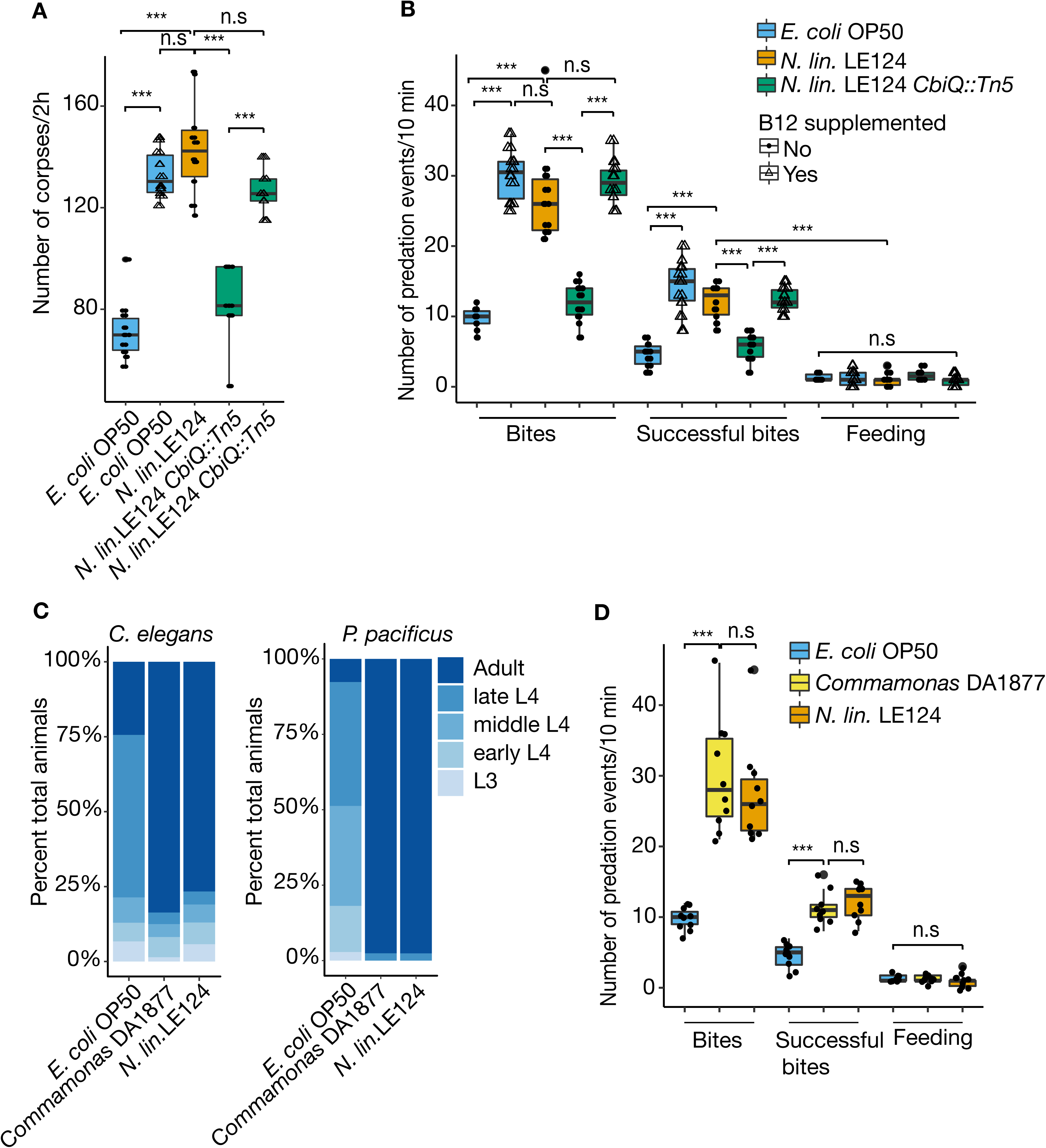
Vitamin B12 containing diet regulates surplus killing behavior and development. (**A**) Corpse assays showing effects of vitamin B12 supplementation on *P. pacificus* predation efficiency with *P. pacificus* fed on either *E. coli* OP50, *N. lin*. LE124, *N. lin*. LE124 *CbiQ::Tn5*, 500nM vitamin B12 supplemented *E. coli* OP50 or 500nM vitamin B12 supplemented *N. lin*. LE124 *CbiQ::Tn5* prior to assays. (**B**) Bite assays showing effects of vitamin B12 supplementation on *P. pacificus* killing behavior with *P. pacificus* fed on either *E. coli* OP50, *N. lin*. LE124, *N. lin*. LE124 *CbiQ::Tn5*, 500nM vitamin B12 supplemented *E. coli* OP50 or 500nM vitamin B12 supplemented *N. lin*. LE124 *CbiQ::Tn5* prior to assays. (**C**) Developmental staging of *C. elegans* and *P. pacificus* showing percentage of L3, early L4, mid L4, late L4 and young adults on plates after feeding with *E. coli* OP50, *Commamonas* DA18877 and *N. lin*. LE124 for either 45 hours (*C. elegans*) or 56 hours (*P. pacificus)*. (**D**) Corpse assays of *P. pacificus* fed with *E. coli* OP50, *Commamonas* DA18877 and *N. lin*. LE124. N=10 replicates for each assay in figure.

Studies by Walhout and co-workers in *C. elegans* showed that developmental acceleration under a *Comamonas aq*. DA1877 diet was also due to vitamin B12^27^. Given the similarities of the *C. elegans* developmental response to *Comamonas* DA1877 and the behavioral response of *P. pacificus* to *N. lin*. LE124, we compared the effect of both bacteria on development and behavior. Indeed, *Comamonas* DA1877 as well as *N. lin*. LE124 induced developmental acceleration of *C. elegans* and *P. pacificus* (Fig. 3C). Similarly, both bacteria enhanced predatory behaviors of *P. pacificus* (Fig. 3D). Thus, the differential effect of bacterial diet on nematode development and behavior might often be due to the uneven distribution of vitamin B12 biosynthesis capabilities of bacteria.

In many animals and humans, vitamin B12 is a co-factor for two enzymes in different pathways (fig. S4A). Methionine-synthase (MS) converts homocysteine to methionine in the cytosolic methionine/S-adenosylmethionine (SAM) cycle and in *C. elegans* is encoded by the *metr-1* gene. The second enzyme, methylmalonyl coenzyme A (CoA) mutase, converts methylmalonyl-CoA to succinyl-CoA in mitochondria and is encoded by the *mce-1* gene in *C. elegans*. In humans, vitamin B12 deficiency causes methylmalonic aciduria and homocysteinemia resulting in devastating diseases^28^. To test if both pathways are required for increased killing behavior in *P. pacificus*, we generated CRISPR/Cas9-derived mutants in *Ppa-metr-1* and *Ppa-mce-1* (fig. S4B, C and D). Both mutants failed to respond to the supplementation of an *E. coli* diet with vitamin B12 (Fig. 4A). Given that SAM is a donor of methyl-groups for many different substrates including RNA, DNA, and proteins, we supplemented an *E. coli* diet of *P. pacificus* wild type and *Ppa-metr-1* mutant animals with methionine. In both cases, methionine supplementation resulted in enhanced killing behavior (Fig. 4B). Thus, both vitamin B12-dependent pathways seem to be involved in *P. pacificus* predatory behaviors.

**Figure 4.**
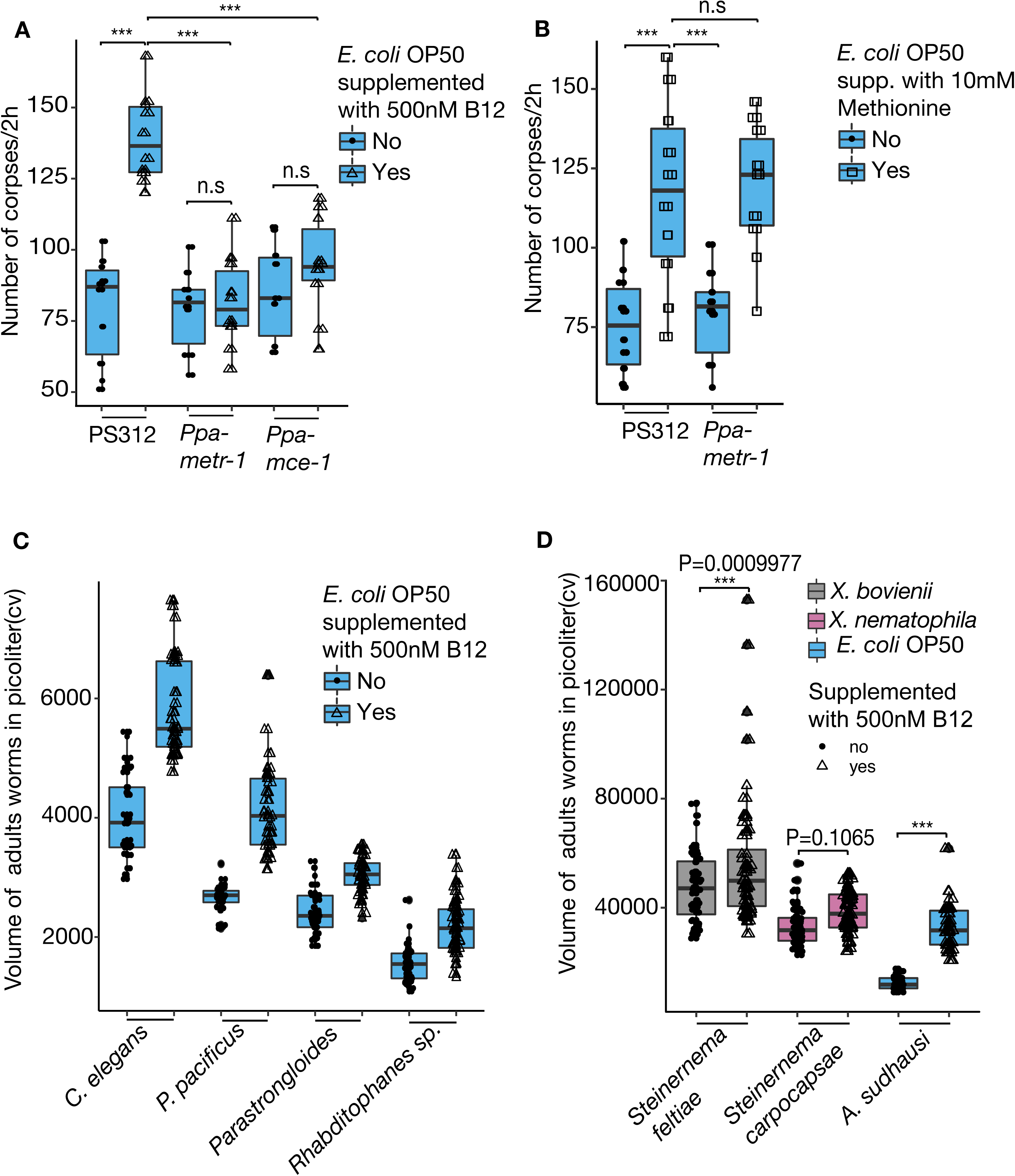
Vitamin B12 influence on development is conserved in various nematodes. (**A**) Corpse assays of *P. pacificus* wild-type (PS312) and mutant animals defective in vitamin B12-dependent pathways *Ppa-metr-1* and *Ppa-mce-1* fed with *E. coli* OP50 supplemented with/without 500nM vitamin B12. (**B**) Corpse assays of PS312 and *Ppa-metr-1* fed with *E. coli* OP50 supplemented with/without 10mM methionine. N=10 replicates for each assay. (**C**) and (**D**) Comparative volume measurement of *C. elegans, P. pacificus, Parastrongyloides trichosuri, Rhabditophanes sp*., *Steinernema carpocapsae and Allodiplogaster sudhausi* after growing on bacterial plates supplemented with vitamin B12 versus non-supplemented plates. N=60 for each assay.

The experiments described above indicate crucial roles of bacterial derived vitamin B12 for the development and behavior of both *P. pacificus* and *C. elegans*. As these nematodes are estimated to have diverged roughly 100 Mya^29^, we next tested how prevalent the effects of vitamin B12 are on the development and physiology of other nematodes, including more distantly related species and representatives that live in diverse ecological settings (supplementary table 1). We grew six nematode species of four major taxonomic clades on a vitamin B12 supplemented diet and measured the effects on their development and growth by quantifying the total worm volume of young adults. In all species tested, we found a significant increase in worm volume (Fig. 4C and D). This included the facultative parasite *Parastrongyloides trichosuri* and the entomopathogenic nematode *Steinernema carpocapsae*. We found the strongest effect on the large free-living nematode *Allodiplogaster sudhausi* that nearly doubled its volume on a vitamin B12 supplemented diet (Fig. 4D). Where possible, we also investigated the effects on developmental speed. Similar to the increase in body size, vitamin B12 supplementation accelerated the development of *Rhabditophanes* sp. and *A. sudhausi* (fig. S4E and F). Taken together, these results demonstrate important physiological and developmental functions of vitamin B12 that are shared across many nematode species.

Our study identified a novel role for nematode-associated microbiota in modulating the complex behavioral trait of predation and therefore, demonstrates a connection between the microbial diet and the nervous system in nematodes. Diverse bacterial species had different effects on the predatory behavioral state with some adversely influencing predation while others enhanced the predatory behaviors. The greatest enhancement in predatory behaviors was observed when *P. pacificus* was fed upon *Novosphingobium* with this increase in killing influenced by bacterial derived vitamin B12. Additionally, we have revealed a more general, conserved role for vitamin B12 in nematode development and growth. Previous studies have shown vitamin B12 to be essential for *C. elegans* development with infertility, growth retardation and a reduction in life-span observed in animals deficient in vitamin B12^27,30,31^. In contrast, behavioral effects have not been reported and similarly, mechanisms of vitamin B12 deficiency in humans that result in neuropathies are currently unknown. It is important to note that the modulation of predation and surplus-killing in *P. pacificus* requires both vitamin B12-dependent pathways. Therefore, we speculate that the influence of vitamin B12 on these behaviors is multifactorial and might well involve several factors. Specifically, the SAM pathway feeds into the methylation of DNA, RNA and proteins, but also lipids and neurotransmitters (fig. S4a). Thus, the presence of vitamin B12 might act through multiple downstream factors, but how it stimulates these effects has yet to be discovered. Importantly however, several neural circuits and neurotransmitter systems of *P. pacificus* have been investigated^6,32-34^. Therefore, future studies can reveal the cellular and molecular foci of vitamin B12-dependence and the influence of the microbiota on nematode predatory behaviors.

## METHODS

### Nematode and Bacterial Strains

A list of all nematode and bacterial species and strains can be found in key resources table.

### Bacterial Culture Conditions

All bacterial strains and mutants were grown overnight in LB (Lysogeny broth) supplemented with 50μg/ml kanamycin where required. Bacteria were grown at 30° g/ml kanamycin where required. Bacteria were grown at 30° C or 37° C depending on the species and 6 cm nematode growth medium (NGM) plates were seeded with 50μg/ml kanamycin where required. Bacteria were grown at 30° l bacterial overnight cultures and were incubated for two days.

### Nematode Culture Conditions

*P. pacificus, C. elegans, Rhabditophanes* sp. KR302 and *A. sudhausi* were grown under standard nematode growth conditions on NGM plates seeded with *Escherichia coli* OP50. Egg cultures were obtained by treating healthy gravid adults with alkaline hypochlorite (bleaching) and were maintained and raised at 20° C on NGM plates. The free-living generation of *Parastrongyloides trichosuri* was cultured as described in Grant et al (2006)^35^. Briefly, to maintain the *P. trichosuri* free-living generation in culture, *E. coli* OP50-spotted NGM plates were incubated for two days at room temperature (RT). Autoclaved rabbit feces were lightly broken and placed on the spotted NGM plate along with *P. trichosuri* animals. Additional *E. coli* OP50 (supplemented with/without vitamin B12) was subsequently added to the dry rabbit feces. The entomopathogenic nematode *Steinernema carpocapsae* was grown on its symbiotic bacterium *Xenorhabdus nematophila*. Symbiotic bacteria were inoculated in LB and incubated at 25°C overnight, 300μg/ml kanamycin where required. Bacteria were grown at 30° l from overnight cultures were spotted to NGM plates (supplemented with/without vitamin B12) and incubated for 1 day at RT. *S._carpocapsae* nematodes were transferred to their respective symbiotic bacterial plates and subsequently grown at 20° C.

### Mouth-form phenotyping

Mouth-form phenotyping was performed as previously reported^6,33^. In brief, axenic worm eggs were obtained by treating healthy gravid *P. pacificus* adults with alkaline hypochlorite, which were subsequently maintained on the test bacteria strains or mutants for at least two generations. Synchronized J4 larvae were picked onto NGM plates with the same test bacteria and roughly 12 hours (hrs) later, worms became young adults. NGM plates with synchronized young adults were placed onto a stereomicroscope with high magnification (150X). The eurystomatous (Eu) mouth form was determined by the presence of a wide mouth, whereas the stenostomatous (St) forms were determined by a narrow mouth. Eu young adult worms were picked for predation assays.

### Predation assays

We used two types of predation assays as described below.

### Corpse assays

Corpse assays facilitated rapid quantification of predatory behavior and were conducted as previously described^6,10,33^. Briefly, in order to generate substantial *C. elegans* larvae for use as prey, cultures were maintained on *E. coli* OP50 bacteria until freshly starved resulting in an abundance of young larvae. These plates were washed with M9 buffer, passed through two Millipore 20μg/ml kanamycin where required. Bacteria were grown at 30° m filters and centrifuged at 377x g to form a concentrated larval pellet. Excess buffer was removed and 1μg/ml kanamycin where required. Bacteria were grown at 30° l of worm pellet was deposited onto a 6 cm NGM unseeded assay plates. This resulted in roughly 3000 prey larvae on each assay plate. Assay plates were left for a minimum of one hour (h) to allow larvae to distribute evenly over the plate. Young adult *P. pacificus* predators were screened for the predatory Eu mouth form and transferred to empty NGM plates for 30 minutes (min) to remove any excess bacteria from their bodies. Subsequently, five *P. pacificus* nematodes were added to each assay plate. Predators were permitted to feed on the prey for two hrs before removal and the plate was subsequently screened for the presence of larval corpses which were identified by the absence of motility coinciding with obvious morphological defects including leaking innards or missing worm fragments. Each assay was replicated ≥5 times. When post-feeding size measurement was required, predatory animals were picked to NGM plates containing no bacteria and measurements were taken using the Wormsizer plug in for Image J/Fiji^36^. See below for Wormsizer experimental details.

### Bite assays

Bite assays provide a more detailed and thorough analysis of the specific interactions associated with predatory behaviors. Bite assays were conducted as previously described^6,10^. Briefly, substantial *C. elegans* prey was generated by maintaining *C. elegans* cultures on *E. coli* OP50 bacteria until freshly starved resulting in an abundance of young larvae. These plates were washed with M9 buffer, passed through two Millipore 20μg/ml kanamycin where required. Bacteria were grown at 30° m filters and centrifuged at 377x g to form a concentrated larval pellet. Excess buffer was removed and 1μg/ml kanamycin where required. Bacteria were grown at 30° l of worm pellet was deposited onto a 6 cm NGM unseeded assay plate. This resulted in roughly 3000 prey larvae on each assay plate. Assay plates were left for a minimum of one h to allow larvae to distribute evenly over the plate. Young adult *P. pacificus* predators were screened for the appropriate predatory Eu mouth morph and transferred to empty NGM plates for 30 min to remove any excess bacteria from their bodies. A single predator was placed on to the assay plate and allowed to recover for 20 min. After recovery, the predatory animal was directly observed under a light stereomicroscope for 10 min and the number of bites, successful bites and feeding events quantified. “Bites” were characterized by a switch to the slower predatory pharyngeal pumping rhythms previously described^6,33^ coinciding with a restriction in movement of the prey. “Successful bites” were characterized by successful rupturing of the prey cuticle resulting in sufficient damage to cause the innards to leak from the wound. “Feeding” was characterized by consumption of the prey through either the observation of prolonged predatory feeding rhythms once the predator had successful grasped its prey, or alternatively, observation of the faster bacterial associated feeding rhythms at the site of a puncture wound. In these assays, no distinction was made as to whether the predatory behavior events were against live prey or against recently killed or wounded animals. Indeed, predators were occasionally observed repeatedly biting the same dying or dead larvae and each contact was quantified as a distinct predatory event. Each assay was conducted with 10 different animals.

### Pharyngeal pumping analysis

*P. pacificus* worms were maintained on 6cm NGM agar plates and fed on the appropriate test bacterial strains prior to assaying. Young adults were transferred onto assay plates and allowed to recover for 15 min from the stress of being transferred. Worms were observed on a Zeiss microscope at 40-63X magnifications, with a high-speed camera and pharyngeal pumping was recorded for 15 seconds, at 50 Hz in at least 20 animals to ensure accurate quantification. The recorded movies were replayed at the desired speed to count individual pumps as previously described^6^.

### *E. coli* OP50 supplementation with *Novosphingobium* L76 supernatant

*E. coli* OP50 and *Novosphingobium* L76 were grown overnight in LB at 37° C and 30° C, respectively. 5ml overnight cultures of each bacteria were grown until they measured an OD_600_ 1. Bacterial cultures were centrifuged at 10000 rpm, RT for 5 min and supernatants were isolated by filtering with 5μg/ml kanamycin where required. Bacteria were grown at 30° m filters. The *E. coli* OP50 pellet was re-suspended with 5ml *Novosphingobium* L76 supernatant. 300μg/ml kanamycin where required. Bacteria were grown at 30° l of the *E. coli* OP50 with *Novosphingobium* L76 supernatant was subsequently spotted to 6 cm NGM plates. OP50 pellet with OP50 supernatant and additionally, *Novosphingobium* L76 were also spotted to 6 cm NGM plates as controls. Spotted NGM plates were ready for assay after two days of incubation. Freshly bleached eggs from well-grown *P. pacificus* cultures were then transferred onto assay plates and worms were transferred to new assay plates two days later. Worms were grown until young adult stage and synchronized young adults were picked and assessed via corpse assays.

### Mixing Bacterial Diets

Liquid cultures of *E. coli* OP50 and *Novosphingobium* L76 were grown in LB at 37° C and 30° C, respectively. Bacterial cultures were diluted to the same OD_600_ and mixed in ratios 1/10,1/100 and 1/1000. Bacterial suspensions were spread onto peptone-free NGM plates to minimize bacterial growth and plates were briefly air dried in a sterile hood. Bleached *P. pacificus* eggs were added to the plates and worms were allowed to grow until young adult stage; synchronized young adults were then picked and assessed via corpse assays.

### Switching bacterial diet

Overnight cultures of *E. coli* OP50 and *Novosphingobium* L76 were spread to NGM plates and incubated at RT for two days. Subsequently, bleached *P. pacificus* eggs were added to the *E. coli* OP50 plates. Worms were transferred from these *E. coli* OP50 plates to *Novosphingobium* L76 at specific developmental stages, L2, L3 and L4, respectively, and were allowed to develop into young adult stage on *Novosphingobium* L76. Worms fed with *E. coli* OP50 or *Novosphingobium* L76 from egg to young adult stage were used as controls. Synchronized young adults were then picked and assessed via corpse assays.

### RNA sequencing

Bacterial strains were grown in LB overnight and spotted to 6 cm NGM plates. Starting from bleached eggs *P. pacificus* nematodes were grown on bacteria for at least two generations and 50 young adults were picked for RNA isolation. Total RNA was extracted using Direct-Zol RNA Mini prep kit (Zymo Research) according to the manufacturer’s guidelines. RNA libraries were prepared by following Truseq RNA library prep kit according to the manufacturer’s guidelines from 1μg/ml kanamycin where required. Bacteria were grown at 30° g of total RNA in each sample (Illumina Company). Libraries were quantified using a combination of Qubit and Bioanalyzer (Agilent Technologies) and normalized to 2.5nM. Samples were subsequently sequenced as 150 bp paired end reads on multiplexed lanes of an Illumina HiSeq3000 (IIlumina Inc). Raw reads have been uploaded to the European Nucleotide archive under the study accession PRJEB33410.

### Analysis of RNA-seq data

The software TopHat (version:2.0.14) was used to align raw reads against the *P. pacificus* reference genome (pristionchus.org, version: Hybrid1) and tests for differential expression were performed by Cuffdiff (version: 2.2.1)^37^. Genes with an FDR-corrected p-value < 0.05 were considered as significantly differentially expressed. For up and downregulated genes, the most significantly enriched metabolic pathways were identified as described previously^12^.

### Generation of transgenic lines

We selected the genes *Ppa-stdh-1* and *Ppa-acs-19.1* to generate transcriptional reporters and established transgenic lines necessary for their use as dietary sensors. For *Ppa-stdh-1*, a 2.3 kb interval encompassing the upstream region and the first two exons was amplified. For *Ppa-acs-19.1*, a 1.4 kb region upstream of the first predicted exon was amplified. These promoters were fused to TurboRFP (Evrogen), together with the 3′ UTR sequence of the gene *Ppa-rpl-23* using the following overlapping primers

*Ppa-stdh-1* - F:

5‘-GCCAAGCTTGCATGCCTGCACATGCTATGGAGCGTAGC-3’;

*Ppa-stdh-1* - R:

5‘-CTGAAAAAAAAAACCCAAGCTTGGGTCCCGAAGACGACGTTGTAGAC-3’;

*Ppa-acs-19.1 -*F

5‘-GGATCCCGTCGACCTGCAGGCATG-3’;

*Ppa-acs-19.1 –*

R 5‘-ATGAGCGAGCTGATCAAG-3’;

*TurboRFP -*F

5‘-TGCATGCCTGCAGGTCGACGGGATCCGCCATCACTATGCATTGCTG-3’ and

*TurboRFP-* R

5‘-TCCTTGATCAGCTCGCTCATCTGAACCAGCAAGGGCGATAG-3’.

PCR fragments were assembled using Gibson assembly kit (NEB) and verified by Sanger sequencing. The *Ppa-stdh-1*::RFP and *Ppa-acs-19.1*::RFP constructs were amplified with the addition of restriction sites (XmaI and PstI) for subsequent digestion. To form stable lines via the formation of complex arrays, the expression construct The *Ppa-stdh-1*::RFP was digested with PstI and 5ng/μg/ml kanamycin where required. Bacteria were grown at 30° l of this, co-injected into the germlines of young adult *P. pacificus* worms with the marker *Ppa-egl- 20*::Venus (10 ng/μg/ml kanamycin where required. Bacteria were grown at 30° l), and genomic carrier DNA (60ng/μg/ml kanamycin where required. Bacteria were grown at 30° l), also digested with PstI^38^. For the *Ppa-acs-19.1*::RFP construct, 10ng/μg/ml kanamycin where required. Bacteria were grown at 30° l of the construct cut with PstI, was injected with the marker *Ppa-egl-20*::RFP (10ng/μg/ml kanamycin where required. Bacteria were grown at 30° l), and genomic carrier DNA (60ng/ μg/ml kanamycin where required. Bacteria were grown at 30° l) also cut with PstI. At least two independent lines were obtained from microinjections for both transgenes.

### Transposon mutagenesis of bacteria

To generate electro-competent cells of *N. lindaniclasticum* LE124 for electroporation, *N. lindaniclasticum* LE124 cells were grown in LB overnight at 30° C. These overnight cultures were diluted (1:10 vol/vol) and incubated for ≅ 6 h to reach early log phase (optical density [OD] at 600nm of 0.3). The culture was centrifuged at 4° C, 10,000 rpm for 10 min before being washed once with ice-cold distillated water and two times with ice-cold 10% glycerol. After the final washing step, cells were centrifuged and the pellet re-suspended with ≅ 1 ml 10% glycerol before 50μg/ml kanamycin where required. Bacteria were grown at 30° l aliquots were distributed to 1.5 ml Eppendorf tubes. The cells in glycerol were electroporated with the EZ-Tn*5* R6Kγ*ori*/KAN-2>Tnp transposon (Epicentre, Madison WI) using an Eppendorf Electroporator 2510 at 2.5 kV yielding around 5 ms. After electroporation, the sample was immediately mixed with SOC (super optimal broth with catabolite repression) medium and incubated at 30° C for two hrs, the culture was then plated on LB agar medium supplemented with 50μg/ml kanamycin where required. Bacteria were grown at 30° g/ml of kanamycin.

### Bacterial transposon mutagenesis library preparation

After two days incubation of the bacteria at 30° C, 10 colonies were randomly selected, picked and a PCR carried out together with Sanger sequencing to confirm the integration of the transposon into the *N. lindaniclasticum* LE124 genome using the primers

KAN-2 FP-1 - F

5‘-ACCTACAACAAAGCTCTCATCAACC-3’ and R6KAN-2 RP-1 - R

5‘-CTACCCTGTGGAACACCTACATCT-3’.

After successful confirmation of the bacterial transposon mutagenesis, around 4500 single mutant colonies were picked and inoculated to 96 well plates in 160μg/ml kanamycin where required. Bacteria were grown at 30° l LB supplemented with 50 μg/ml kanamycin where required. Bacteria were grown at 30° g/ml of kanamycin. Overnight cultures of all mutants were mixed with 160μg/ml kanamycin where required. Bacteria were grown at 30° l 50% glycerol and frozen at −80°C.

### Transposon mutant library screening using dietary sensors

Transposon mutants were inoculated into 96 well plates in 180μg/ml kanamycin where required. Bacteria were grown at 30° l LB supplemented with 50μg/ml kanamycin where required. Bacteria were grown at 30° g/ml of kanamycin. After overnight growth at 30° C, 20μg/ml kanamycin where required. Bacteria were grown at 30° l from the mutant cultures were spotted to 24-well NGM plates. Bacterial mutant strains were incubated for two days and eggs of *P. pacificus* RS3271 (*Ppa-stdh-1::*RFP) or *P. pacificus* RS3379 (*Ppa-acs-19.1*::RFP) were bleached and filtered with Millipore 120.0μg/ml kanamycin where required. Bacteria were grown at 30° m filters to reduce the amount of adult worm carcasses. Around 50-100 bleached eggs were spotted to each well with mutant bacteria; *E. coli OP50* and *N. lindaniclasticum* LE124 wild type strain were used as controls. Fluorescent worms were grown on the bacterial strains until they became young adults. The *Ppa-stdh-1::*RFP line was screened for decreased RFP expression while the *Ppa-acs-19.1*::RFP line was screened for increased RFP expression. Initial positive results were re-screened at least three times to confirm changes in gene expression.

### Analysis of Transposon Mutant Sequencing Data

Raw reads were aligned against *N. lindaniclasticum* LE124 reference genome and transposon sequence by the BWA aln and samse programs (version 0.7.12-r1039)^39^. The generated sam files were screened for read pairs where one read aligned to the transposon sequence and the second read was unmapped. The location of the affected gene was identified by realignment of the unmapped second read against the *N. lindaniclasticum* LE124 reference with the help of blastn (version: 2.6.0)^40^.

### Generation of CRISPR-induced mutants of *Ppa-metr-1* and *Ppa-mce-1*

We generated mutant alleles for *Ppa-metr-1* and *Ppa-mce-1* using the CRISPR/Cas9 technique following the protocol described previously (Witte et al, 2015). crRNAs were synthesized by Integrated DNA Technologies and fused to tracrRNA (also Integrated DNA Technologies) at 95° C for five min before the addition of the Cas9 endonuclease (New England Biolab). After a further five min incubation at RT, TE buffer was added to a final concentration of 18.1μg/ml kanamycin where required. Bacteria were grown at 30° M for the sgRNA and 2.5μg/ml kanamycin where required. Bacteria were grown at 30° M for Cas9. Around 20 young adults were injected; eggs from injected P0s were recovered up to 16 hrs post injection. After hatching and two days of growth these F1 were picked onto individual plates until they had also developed and laid eggs. The genotype of the F1 animals was subsequently analyzed via Sanger sequencing and mutations identified and isolated in homozygosity.

### Phylogenetic Analysis

For two fatty acid metabolism related genes with differential expression between the bacterial diets, we retrieved homologs by BLASTP searches against WormBase (version: WS270) and pristionchus.org (version: TAU2011). Homologous protein sequences from *C. elegans* and *P. pacificus* were aligned by MUSCLE (version: 3.8.31)^41^) and maximum likelihood trees were generated with the help of the phangorn package in R (version: 3.5.3, parameters: model=“LG”, optNni=TRUE, optBf=TRUE, optInv=TRUE)^42^. To assess the robustness of the resulting trees, 100 bootstrap pseudoreplicates were calculated. For two *C. elegans* candidate genes involved in the Vitamin B12 pathway, one-to-one orthologs in *P. pacificus* could directly be retrieved from BLASTP searches against WormBase (version: WS270): *Ppa-metr-1* (PPA25255) and *Ppa-mce-1* (PPA39850). One-to-one orthology was confirmed by phylogenetic analysis.

### Metabolite supplementation

Methylcobalamin (Vitamin B12 CAS Number 13422-55-4) and L-methionine (CAS Number 63-68-3) were purchased from Sigma and dissolved in water at the highest possible soluble concentrations to prepare stock solution. A methylcobalamin stock was prepared fresh before use in each experiment. Metabolite solutions were mixed with NGM agar at the required concentration just before pouring the 6 cm plates. Plates were allowed to dry at RT for two days and then spotted with *E. coli* OP50.

### *Ppa*-*acs-19*.*1*::RFP gene expression screening on metabolite supplemented plates

We used *Ppa*-*acs-19.1*::RFP transgenic animals to determine working concentrations of metabolite supplementations. Bleached *Ppa*-*acs-19.1*::RFP transgenic eggs were transferred to metabolite-supplemented plates, which were prepared as described above. *Ppa*-*acs-19.1*::RFP positive young adults were screened for differences in gene expression in comparison to control animals grown on a *E. coli* OP50 and *N. lindaniclasticum* LE124 diet without metabolite supplementation.

### Imaging transgenic reporter lines

Eggs of transgenic reporter lines *Ppa*-*acs-19.1*::RFP and *Ppa-stdh-1*::RFP were bleached and transferred to bacteria plates that were prepared as described. Three ml of 2% agar was prepared and a drop (150μg/ml kanamycin where required. Bacteria were grown at 30° l) of 1 M sodium azide (NaN_3_) was added and mixed with agar to immobilize the worms. Around 200-μg/ml kanamycin where required. Bacteria were grown at 30° l agar was dropped on microscope slide and young adult transgenic worms were placed on the agar. Images of the worms were taken with 10X objective of ZEISS Imager Z1 equipped with the AxioCam camera using ZEN imaging software. The same exposure time was applied to all images.

### Vitamin B12 (Methylcobalamin) supplementation assays

Vitamin B12-supplemented plates were prepared as described above. *P. pacificus, C. elegans, Rhabditophanes* sp. KR3021, *A. sudhausi* SB413, as well as *Ppa-metr-1* (*tu1436, tu1436*) and *Ppa-mce-1* (*tu1433, tu1434* and *tu1435*) mutant animals were grown on supplemented plates from egg to young adult stage and subsequently used for i) predatory assays, ii) worm size measurements and iii) developmental assays. For supplementation experiments with free-living *P. trichosuri*, J2 larvae were washed five times with M9 medium and filtered with Millipore 20.0μg/ml kanamycin where required. Bacteria were grown at 30° m filters before being soaked in PBS supplemented with 100μg/ml kanamycin where required. Bacteria were grown at 30° g/ml penicillin and ampicillin for one h to avoid contamination. J2 larvae were washed a final time with PBS containing no antibiotics and transferred to assay plates. For *S. carpocapsae*, J2 larvae were washed with M9 medium and filtered with Millipore 20.0μg/ml kanamycin where required. Bacteria were grown at 30° m filters before transferring to NGM plates supplemented with/without 500nM vitamin B12.

### Worm size measurement

*P. pacificus, C. elegans, Rhabditophanes* sp., *P. trichosuri, A. sudhausi* and *S. carpocapsae* synchronized young adults were transferred from assay plates to NGM plates without bacteria. Bright field images of the worms were taken using 0.63x objective of ZEISS SteREO Discovery V12 using the AxioCam camera. Images were analyzed using the Wormsizer plug in for Image J/Fiji^36^. Wormsizer detects and measures the volume of the worms.

### Development rate assays

For development rate assays, *P. pacificus, C. elegans, Rhabditophanes sp*. and *A. sudhausi* were grown on OP50 at 20° C. Nematode eggs were bleached, washed with M9 several times and allowed to hatch in M9 medium for 20 hrs in the absence of food to cause J2 arrest. Once synchronized, J2 larvae were filtered through two Millipore 20.0μg/ml kanamycin where required. Bacteria were grown at 30° m filters and around 30-60 J2 animals were transferred to NGM plates (supplemented with/without 500nM vitamin B12) spotted in 50μg/ml kanamycin where required. Bacteria were grown at 30° l of the desired test bacterial strain. Nematodes were subsequently allowed to develop on test bacteria for the following time periods: *P. pacificus 57* hrs at 20° C, *C. elegans* and *Rhabditophanes sp*. 45 hrs at 20° C and *A. sudhausi* for 144 hrs at RT. Following this, worms were categorized into groups based on the development of the vulva and germ line using 0.63x objective of ZEISS SteREO Discovery V12 following previously established protocols^27^.

### Statistical analysis

Statistical calculations (mean, SEM, and t test) were performed by using R studio software. Pairwise t-tests with Benjamini-Hochberg multiple testing correction were applied when testing the effect of a single treatment or mutant against one single control sample. For tests across different groups (e.g. treatments, mutants, behaviors), Tukey-HSD test was applied. Significance is designated between two samples according to the following scale: 0 ‘***’ 0.001 ‘**’ 0.01 ‘*’ 0.05 ‘n.s’ 0.1 ‘n.s’ 1.

## ACKNOWLEDGMENTS

We thank Dr. A. Streit and R. Ehlers for *Parastrongyloides* and *Steinernema* material, respectively, and members of the Sommer lab for discussion. This work was funded by the Max Planck Society.

## AUTHOR CONTRIBUTIONS

N.A. and J.W.L. performed all behavioral experiments. W.R. performed the RNA-seq experiments, H.W., N.A. and J.W.L. generated dietary sensor lines and CRISPR-induced mutants. Bioinformatic analysis was performed by W-S.L. and C.R. All experiments were designed by N.A., C.R., J.W.L. and R.J.S.

## DATA AND MATERIAL AVAILABILITY

RNA-seq data has been deposited at the European Nucleotide Archive under the study accession PRJEB33410. All other data is available in the main text or the supplementary materials.

## SUPPLEMENTARY FIGURE LEGENDS

**Figure S1.**
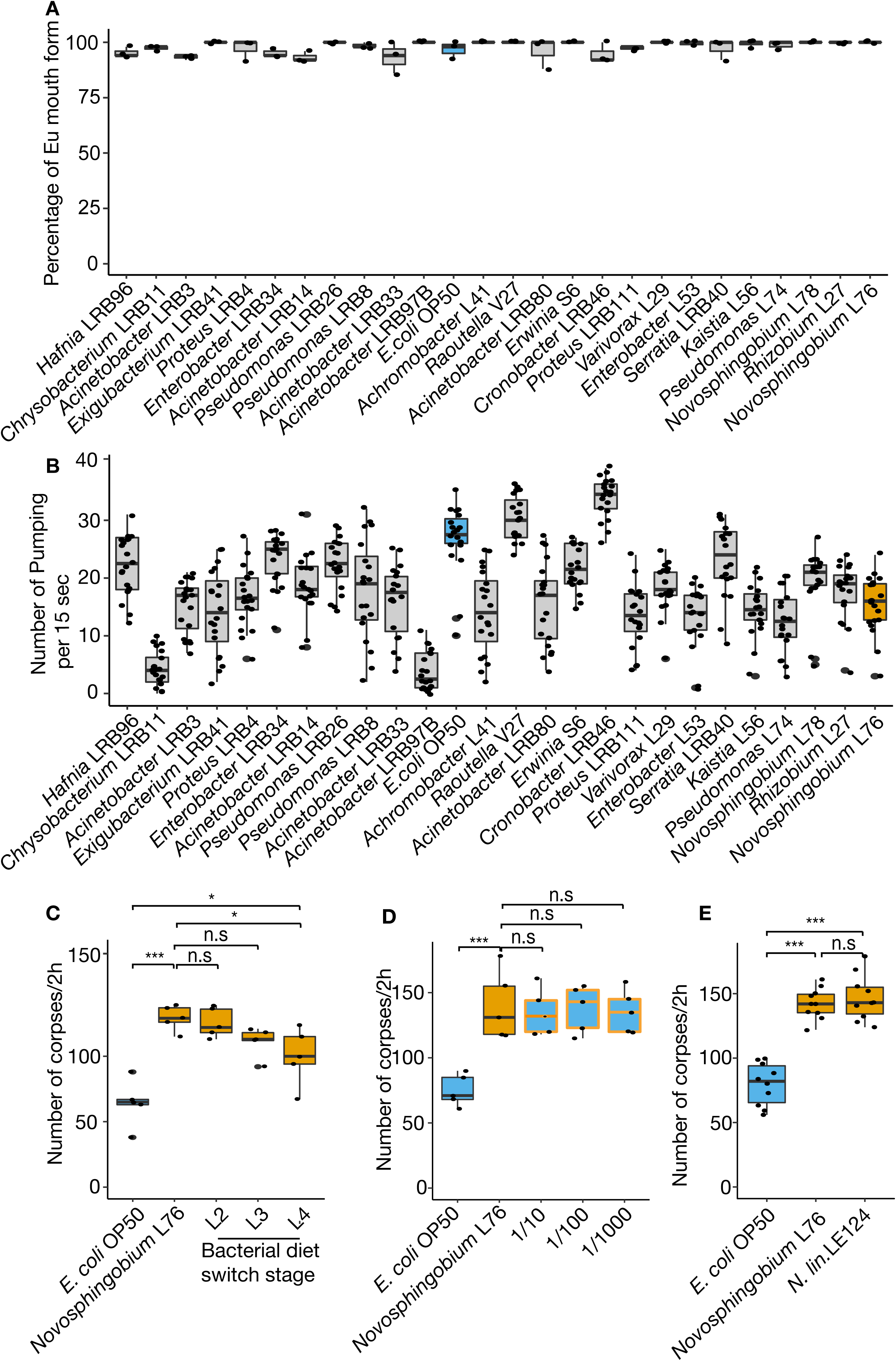
Bacterial diet affects predatory behavior in *P. pacificus*. (**A**) Mouth form ratio of *P. pacificus* PS312 after feeding with 25 different bacteria strains. Bacterial diet fails to influence mouth-form ratio. N=3 replicates for each assay. (**B**) Pharyngeal pumping behavior of *P. pacificus* PS312 on 25 different bacterial diets. N=20 replicates for each assay. (**C**) Corpse assay illustrating affect of bacterial diet switching from *E. coli* OP50 to *Novosphingobium* L76 at particular *P. pacificus* development stages. Corpse assays were performed with young adults suggesting feeding with *Novosphingobium* L76 at diverse developmental stages modify killing behavior. (**D**) Corpse assays of *P. pacificus* previously fed with a mixture of *Novosphingobium* L76 and *E. coli* OP50 at 1/10, 1/100 and 1/1000 concentrations. Low concentrations of *Novosphingobium* L76 in the diet is sufficient to influence killing behavior. Bacteria were spotted to NGM without peptone to prevent bacterial growth. N=10 replicates for each assay. **(E)** Corpse assays of *P. pacificus* previously fed on either Novosphingobium L76 or Novosphingobium LE124. The increased killing behaviors are observed in both strains of Novosphingobium. N=10 replicates for each assay.

**Figure S2.**
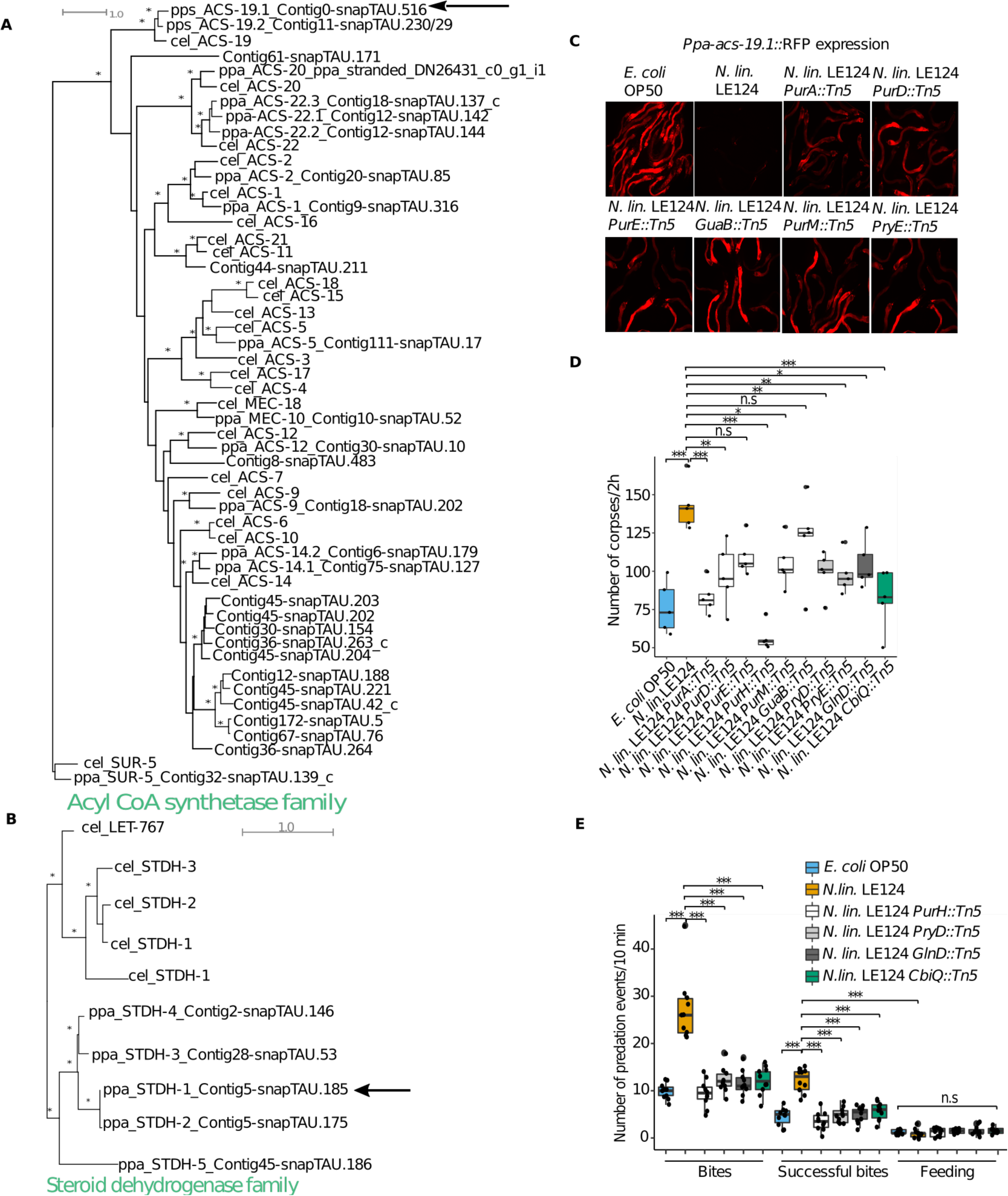
Mutations in multiple pathways affect dietary sensor expression and predatory behavior. **(A)** A phylogenetic analysis of acs-19 and let-767 homologs indicates that individual members of the Acyl CoA synthtase family and **(B)** the steroid dehydrogenase family (panel B) have undergone lineage specific duplications. Nodes with bootstrap support ≥ 90/100 are labeled with stars and arrows mark *P. pacificus* genes that were used as dietary sensors. (**C**) Images of *Ppa-acs-19.1*::RFP dietary sensor showing purine (*N. lin*. LE124 *PurA::Tn5, N. lin*. LE124 *PurD::Tn5, N. lin*. LE124 *PurE::Tn5, N. lin*. LE124 *GuaB::Tn5* and *N. lin*. LE124 *PurM::Tn5)* and pyrimidine biosynthesis (*N. lin*. LE124 *PryE::Tn5*) mutants increase the expression of the dietary sensor in comparison to *N. lin*. LE124 wild-type diet. (**D**) Corpse assays of *P. pacificus* fed with *N. lin*. LE124 mutants from vitamin B12 (green), purine (white), pyrimidine biosynthesis (grey) and nitrogen metabolism (dark grey) all decreasing killing efficiency in comparison to *N. lin*. LE124 wild-type diet. N=10 replicates for each assay. (**E**) Bite assays of *P. pacificus* previously fed on *E. coli* OP50, *N. lin*. LE124 and *N. lin*. LE124 mutants from vitamin B12 (green), purine (white), pyrimidine biosynthesis (grey) and nitrogen metabolism (dark grey) modulating killing efficiency. Ten replicates for each assay.

**Figure S3.**
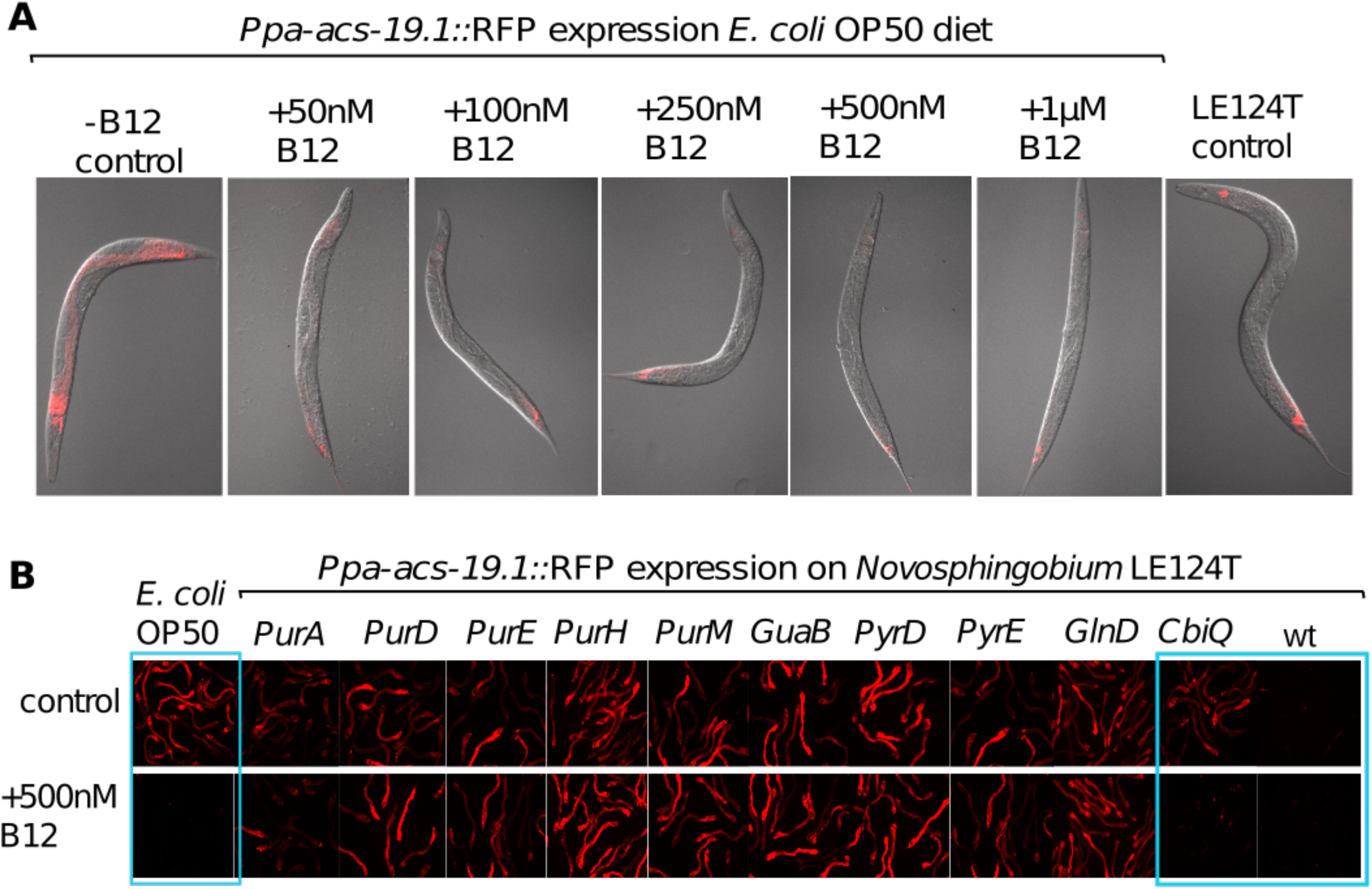
Vitamin B12 regulates fatty acid gene expression and development. **(A)** *Ppa-acs-19.1* transgenic worms were grown on NGM plates supplemented with various concentrations of vitamin B12. NGM plates without vitamin B12 spotted with *E. coli* OP50 and *N. lin*. LE124 were used as controls. Images of transgenic animals were taken to determine the most efficient vitamin B12 concentration. Vitamin B12 Supplemented *E. coli* OP50 phenocopies *N. lin*. LE124 effect on *Ppa-acs-19.1* expression. **(B)***Ppa-acs-19.1* transgenic worms were added to NGM plates with *N. lin*. LE124 transposon mutants and with/without supplementation with 500 nM vitamin B12. *E. coli* OP50 and *N. lin*. LE124 were as controls. Vitamin B12 supplementation rescued *Ppa-acs-19.1* expression on *N. lin*. LE124 *CbiQ::Tn5* mutant (blue highlighted box).

**Figure S4.**
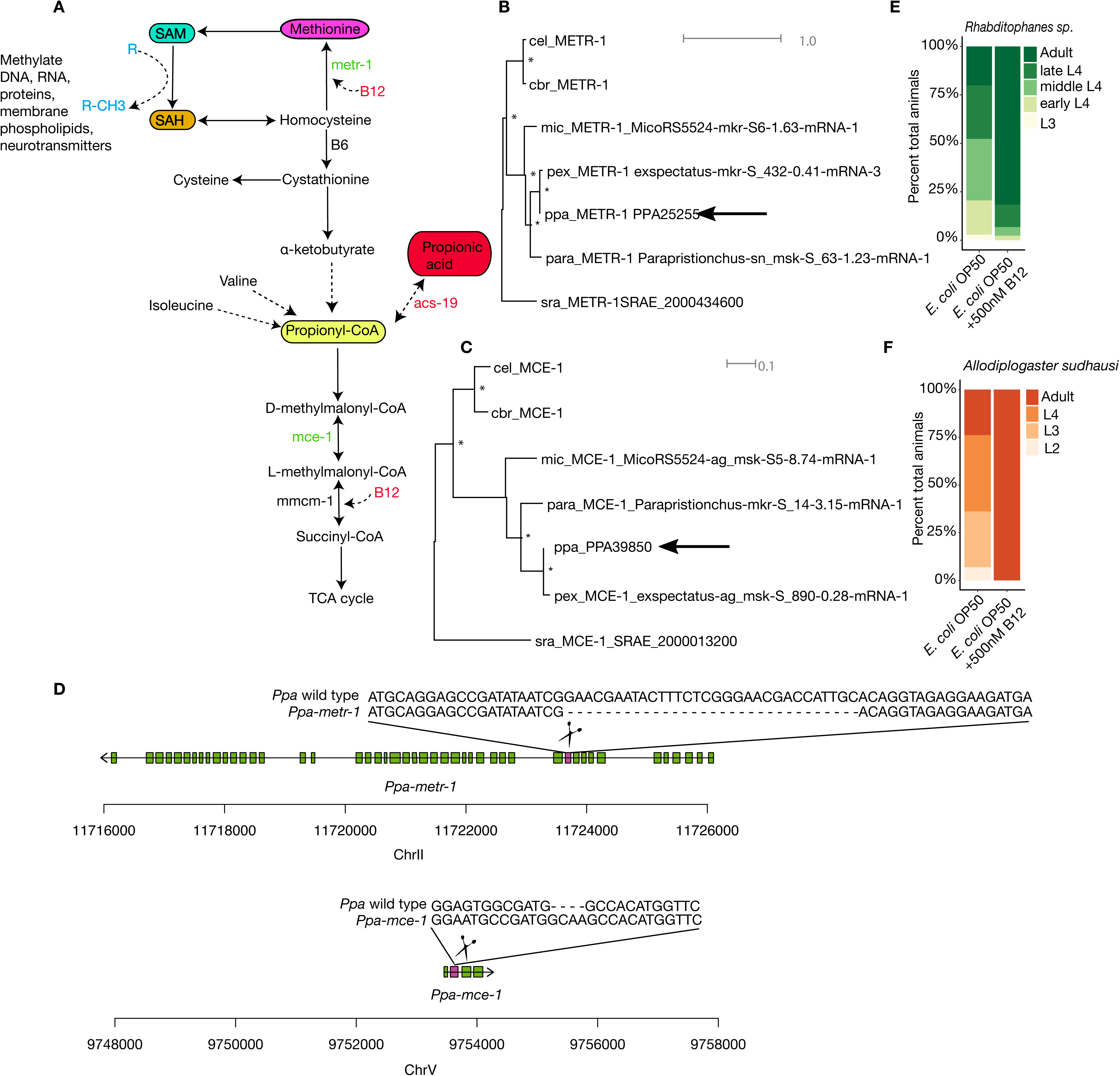
Vitamin B12 dependent metabolic pathways. **(A)** Network of the main two vitamin B12-dependent pathways. *P.pacificus* Orthologous of genes labeled in green were mutated with CRISPR/Cas9. Orthologous of red-labeled *acs-19* used as dietary sensor. **(B)** One-to-one orthologs could be identified for *metr-1* **(C)** and *mce-1*. Nodes with bootstrap support ≥ 90/100 are labeled with stars and arrows mark *P. pacificus* genes that were used for functional studies. **(D)** Mutations were induced in both *Ppa- metr-1* and *Ppa-mce-1* using CRISPR/Cas9 with the target locations indicated in both genes (scissors). Mutations induced via CRIPSR/Cas9 are also shown. (**E**) and (**F**) Developmental staging of *Rhabditophanes sp*. and *A. sudhausi* on *E. coli* OP50 NGM plates supplemented with/without vitamin B12. The development of *Rhabditophanes sp*. and *A. sudhausi* was accelerated with vitamin B12 supplementation. N=10 replicates for each assay.

## SUPPLEMENTARY TABLE LEGENDS

**Table S1**. List of strains and other resources that were used in this study.

**Table S2**. List of differentially expressed genes between *P. pacificus* grown on *E.coli* OP50 and *Novosphingobium* L76. List includes *P. pacificus* gene identifiers, the associated expression fold changes, FDR corrected P-values and where appropriate the identified *C. elegans* orthologous genes can be found in a separate excel file.

---

**Supplementary Table 1.**
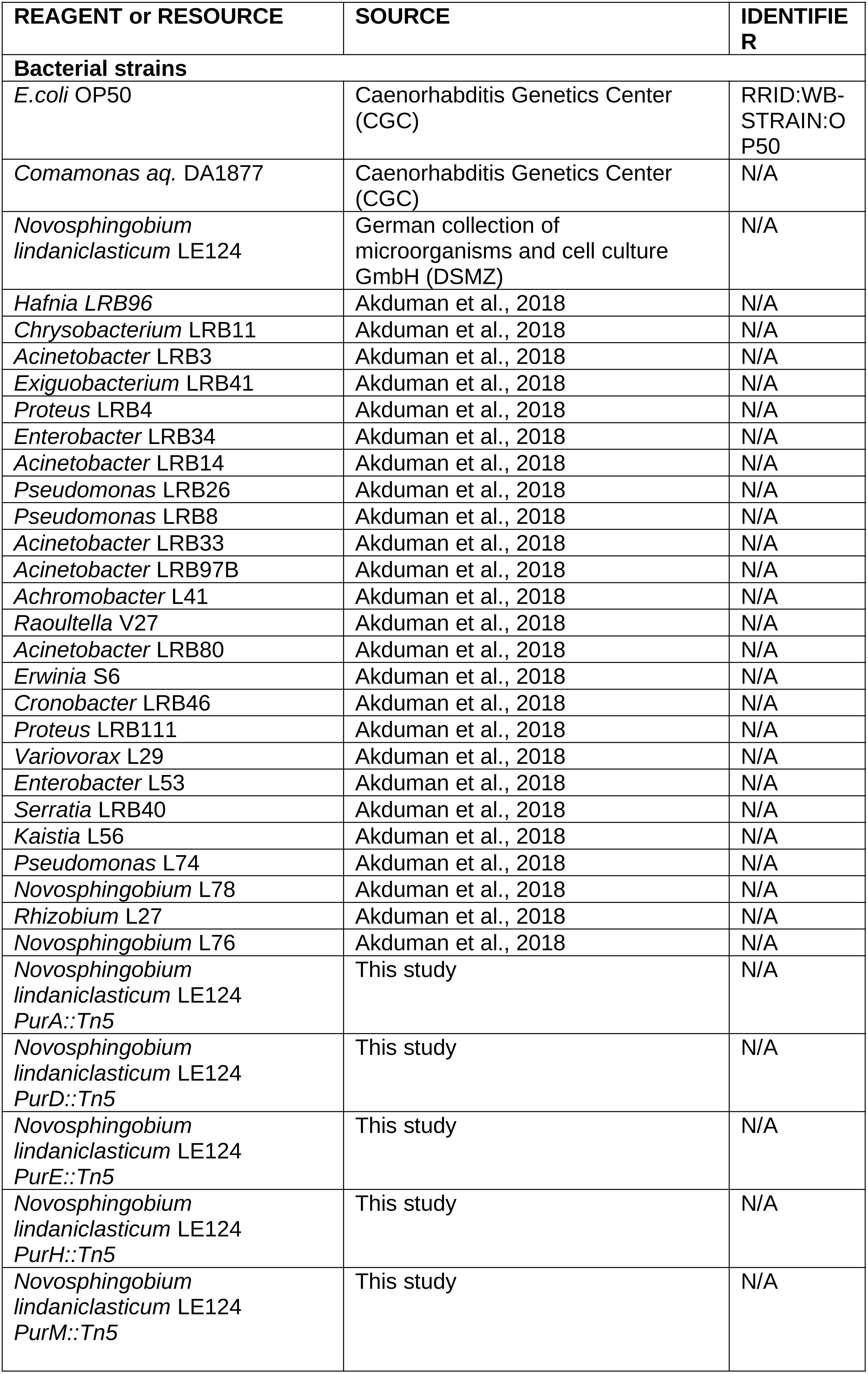
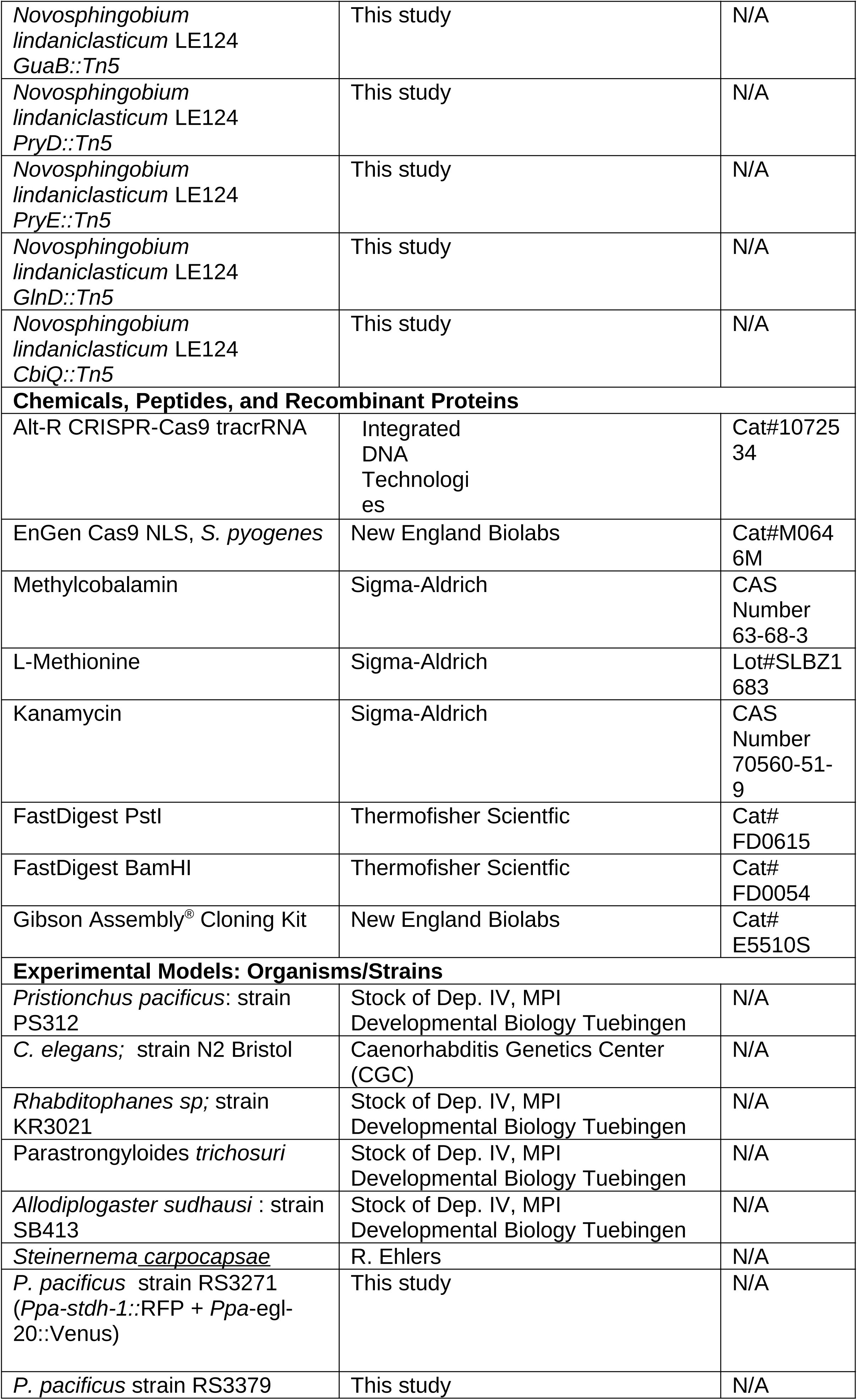
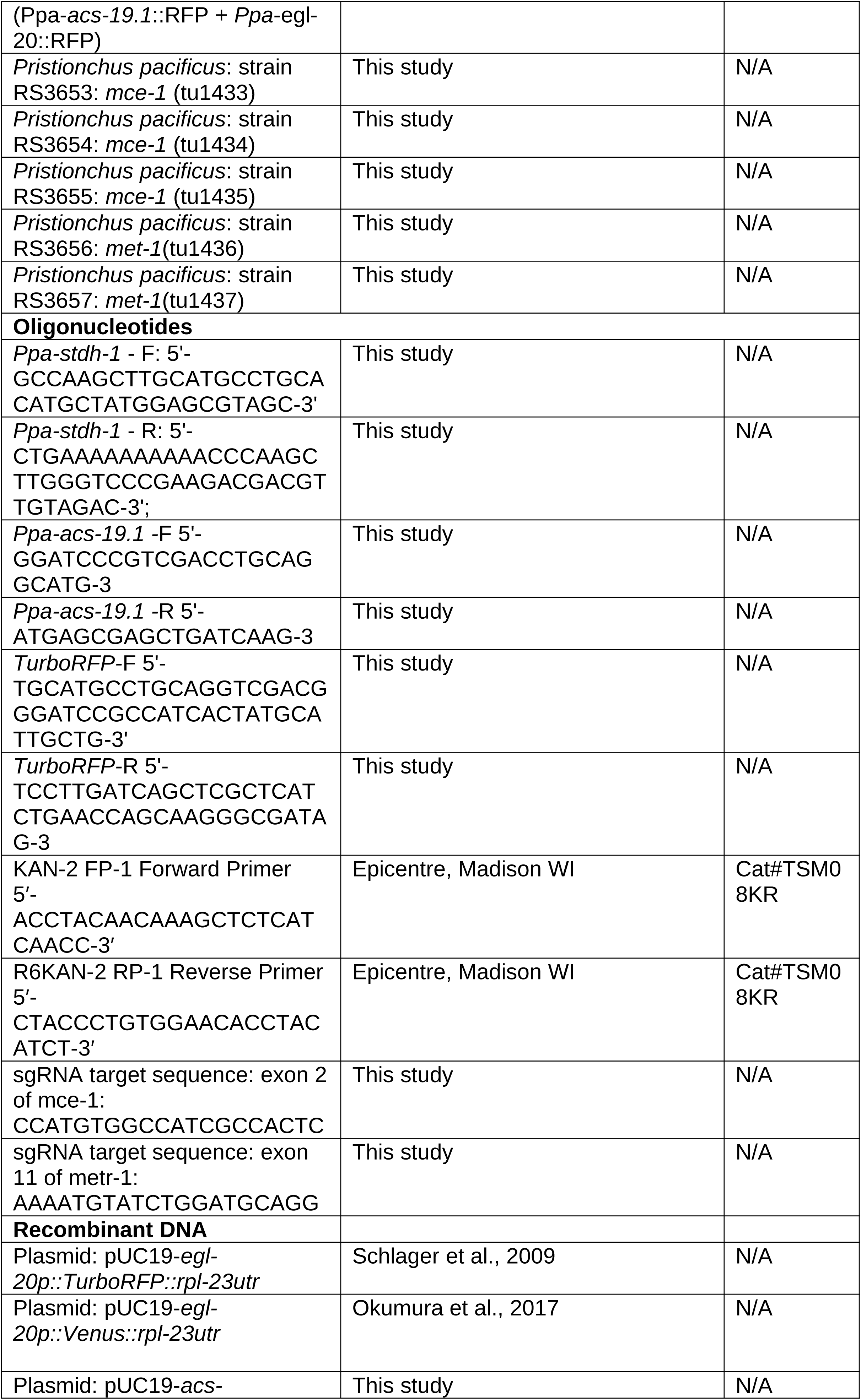
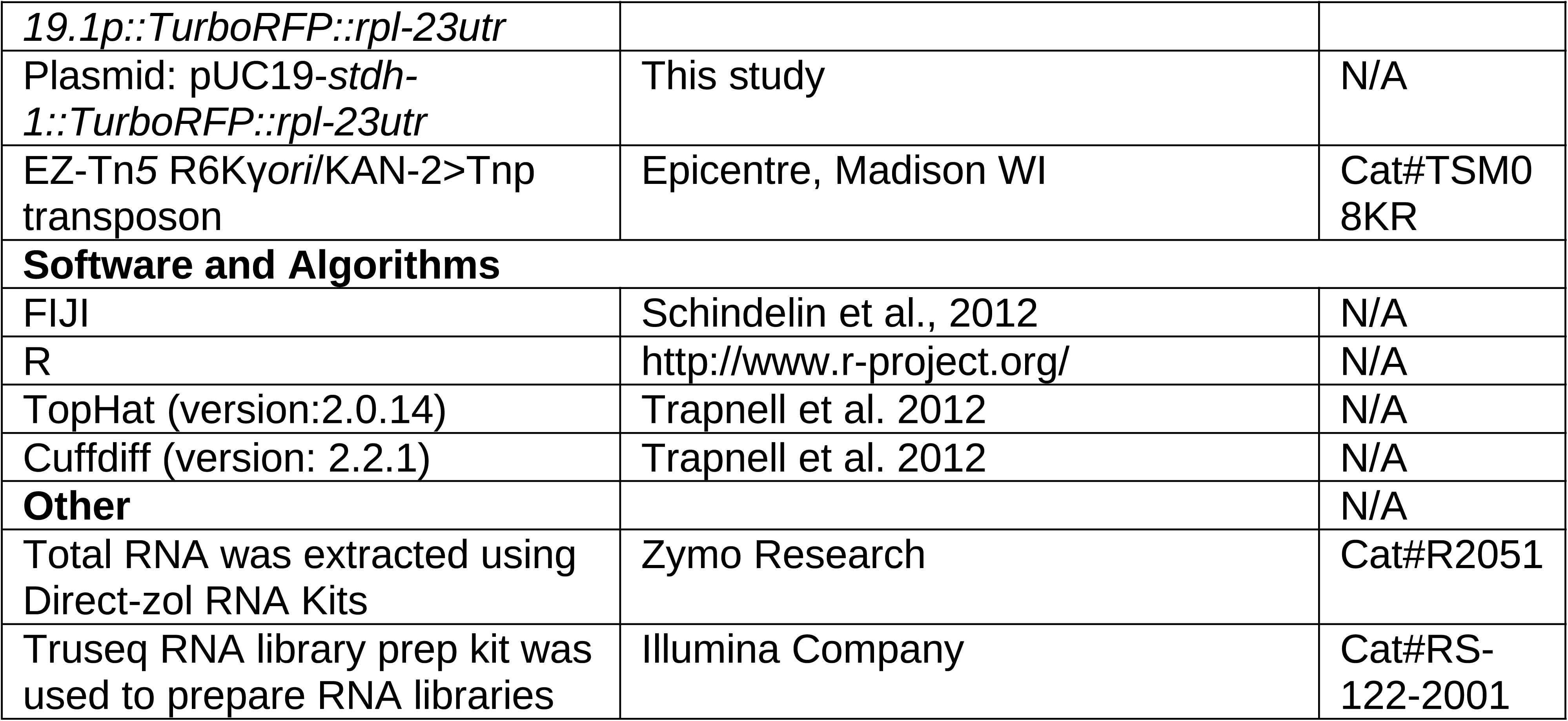

